# Protein Domain Hotspots Reveal Functional Mutations across Genes in Cancer

**DOI:** 10.1101/015719

**Authors:** Martin L. Miller, Ed Reznik, Nicholas P. Gauthier, Bülent Arman Aksoy, Anil Korkut, Jianjion Gao, Giovanni Ciriello, Nikolaus Schultz, Chris Sander

## Abstract

In cancer genomics, frequent recurrence of mutations in independent tumor samples is a strong indication of functional impact. However, rare functional mutations can escape detection by recurrence analysis for lack of statistical power. We address this problem by extending the notion of recurrence of mutations from single genes to gene families that share homologous protein domains. In addition to lowering the threshold of detection, this sharpens the functional interpretation of the impact of mutations, as protein domains more succinctly embody function than entire genes. Mapping mutations in 22 different tumor types to equivalent positions in multiple sequence alignments of protein domains, we confirm well-known functional mutation hotspots and make two types of discoveries: 1) identification and functional interpretation of uncharacterized rare variants in one gene that are equivalent to well-characterized mutations in canonical cancer genes, such as uncharacterized *ERBB4* (S303F) mutations that are analogous to canonical *ERRB2* (S310F) mutations in the furin-like domain, and 2) detection of previously unknown mutation hotspots with novel functional implications. With the rapid expansion of cancer genomics projects, protein domain hotspot analysis is likely to provide many more leads linking mutations in proteins to the cancer phenotype.

## INTRODUCTION

The landscape of somatic mutations in cancer is extraordinarily complex, making it difficult to distinguish oncogenic alterations from passenger mutations. Many approaches use the recurrence of alterations in a single gene across tumor samples to identify potential driver genes. However, the molecular functions of genes are often pleiotropic, and in many cases it may not be a gene as an entity itself, but rather the specific function of a gene or a set of genes that is under selective pressure in cancer. For example, in T-cell acute lymphoblastic leukemia, the transmembrane signaling receptor *NOTCH1* is activated by mutations in the heterodimerization and the PEST domains (Weng *et al*., 2004), while in squamous cell carcinomas, notch-signaling has a tumor suppressive role and notch receptors (*NOTCH1-4*) are inactivated by mutations in the ligand binding EGF-like domains (Wang *et al*., 2011). Thus, an alternative approach to assessing the relevance of somatic alterations is to determine the recurrence of mutations in genes involved in similar molecular functions. One powerful method for systematically assessing common biological function of genes is through the analysis of protein domains, which are evolutionarily conserved, structurally related functional units encoded in the protein sequence of genes (Holm & Sander, 1996; Chothia *et al*., 2003). By coupling the observation of mutations across genes in a domain family together, it may be possible to identify putative functional alterations that confer a selective, functional advantage to cancer cells.

Large cross-institutional projects, such as The Cancer Genome Atlas (TCGA), have recently profiled the major human cancer types genomically, including glioblastoma (McLendon *et al*., 2008), lung (Hammerman *et al*., 2012; Ding *et al*., 2008), ovarian (Bell *et al*., 2011), breast (Koboldt *et al*., 2012), endometrial (Getz *et al*., 2013), kidney (Creighton *et al*., 2013) and colorectal cancer (Cancer Genome Atlas Network, 2012). Through whole-exome sequencing (WES) of tumor-normal pairs, these and other studies have provided catalogues of somatically mutated genes that are frequently altered and therefore likely associated with disease development. However, despite a collection of mutation data from nearly 5,000 samples encompassing 21 tumor types, the results from a recent pan-cancer study illustrate that by using recurrence of mutations in genes, thousands of samples per tumor type are needed to confidently identify genes that are mutated at low but clinically relevant frequencies (2-5%) (Lawrence *et al*., 2014a).

Several analytical approaches have been developed to detect genes associated with oncogenesis (Gonzalez-Perez & Lopez-Bigas, 2012; Dees *et al*., 2012; Lawrence *et al*., 2014b). One of these widely applied algorithms, MutSigCV, compares the gene-specific mutation burden to a background model using silent mutations in the gene and gene neighborhood to estimate the probability that the gene is significantly mutated (Lawrence *et al*., 2014b). Importantly, the method also incorporates contextual information, such as genomic parameters that correlate strongly with the mutation background rate (DNA replication timing and the general level of transcriptional activity) (Lawrence *et al*., 2014b) as well as the tendencies of mutations to cluster to specific sites and to occur at positions that are evolutionarily conserved (Lohr *et al*., 2012; Lawrence *et al*., 2014a). Additional approaches have been developed to predict the functional impact of specific amino acid changes. These approaches generally rely on analyzing physicochemical properties of amino acid substitutions (*e.g.*, changes in size and polarity), structural information (*e.g.*, hydrophobic propensity and surface accessibility), and the evolutionary conservation of the mutated residues across a set of related genes (Reva *et al*., 2011; Yue *et al*., 2006; Bromberg *et al*., 2008; Ng, 2003; Adzhubei *et al*., 2010). Other approaches analyze mutations across sets of functionally related genes (*e.g.*, genes in the same signaling pathway) to test for a possible enrichment of mutation events (Cerami *et al*., 2010; Ciriello *et al*., 2012; Hofree *et al*., 2013; Torkamani & Schork, 2009).

Protein domains represent particular sequence variants that have been formed over evolution by duplication and/or recombination (Holm & Sander, 1996; Chothia *et al*., 2003). Domains often encode structural units associated with specific cellular tasks, and large proteins with multiple domains can have several molecular functions each exerted by a specific domain. The structure-function relationship encoded in domains has been used as a tool for understanding the effect of mutations across functionally related genes. For example, some of the most frequent oncogenic mutations in human cancer affect analogous residues of the activation segment of the kinase domain and cause constitutive activation of several oncogenes, including FLT3 D835 mutations in acute myeloid leukemia, *KIT* D816 mutations in gastrointestinal stromal tumors, and *BRAF* V600 mutations in melanoma (Dibb *et al*., 2004; Greenman *et al*., 2007). In the *SMAD* tumor suppressor genes, mutations in conserved residues of the MAD homology 2 (MH2) domain have analogous effects in *SMAD2* and *SMAD4*, disrupting homo- and hetero-oligomeric interactions critical for SMAD signaling (Shi *et al*., 1997). Proteome-wide bioinformatics analysis of mutations in domains have been performed to identify domains enriched for alterations (Nehrt *et al*., 2012; Peterson *et al*., 2012) as well as to detect significantly mutated domain hotspots through multiple sequence analysis (Peterson *et al*., 2010; Yue *et al*., 2010). However, due in part to the scarcity of data available at the time of analysis, these studies did not perform a systematic pan-cancer analysis of mutations in domains and provided limited biological insights.

Here, we performed a systematic and comprehensive analysis of mutations in protein domains using data from more than 5,000 tumor-normal pairs from 22 cancer types profiled by the TCGA consortium and domains from the protein family database Pfam-A (Punta *et al*., 2011). We confirmed that signaling domains in canonical oncogenes are recurrently altered in cancer and further identified domains that are enriched for mutations contributed by infrequently altered genes not previously associated with cancer. Using multiple sequence analysis, we determined if conserved residues in protein domains were affected by mutations across related genes. This analysis enabled us to identify putative “domain hotspots”. For example, we discovered novel hotspots with putative driver mutations in the prolyl isomerase domain and in the DNA-binding forkhead domain. We further exposed rare mutations that associated with well-characterized oncogenic mutations, including the furin-like domain where uncharacterized mutations in *ERBB4* (S303 F) are analogous to known oncogenic mutations in the same domain of *ERRB2* (S310 F), suggesting similar functional consequences. In several cases, we associated rare mutations in potential cancer genes with therapeutically actionable hotspots in known oncogenes, underlining the potential clinical implications of our findings. We have made all results freely available to the research community through an interactive web resource (http://www.mutationaligner.org) that will be continuously updated as data become available from cancer genomics projects.

## RESULTS

### Mapping somatic mutations to protein domains

To systematically analyze somatic mutations in the context of conserved protein domains, we collected WES data from 5496 tumor-normal pairs of 22 different tumor types profiled by the TCGA consortium. To obtain a uniform data set of mutation calls, annotation of somatic mutations were based on the publicly available data (Oct 2014) from the cBioPortal for cancer genomics data (Cerami *et al*., 2012; Gao *et al*., 2013) (Fig. 1). After filtering out ultra-mutated samples and mutations in genes with low mRNA expression levels (**Methods**), the data consisted of a total of 727,567 mutations in coding regions with 463,842 missense, 192,518 silent, and 71,207 truncating or small inframe mutations (**Supplementary Fig. 1**). Focusing on missense mutations, we observed that the relative proportion of amino acids affected by mutations varied considerably between cancer types (Fig. 2). These amino acid mutation biases are due to a combination of variations in the codon usage between different amino acids and the variations in the base-pair transitions and transversions observed between different cancer types (Lawrence *et al*., 2014b; Alexandrov *et al*., 2013). Because of the high mutation rate of CG dinucleotides across all cancers, arginine (R) is the most frequently altered amino acid despite being the 9*^th^* most common amino acid as CG dinucleotides are present in four out of six of arginine’s codons (**Supplementary Fig. 2**)

**Figure 1:**
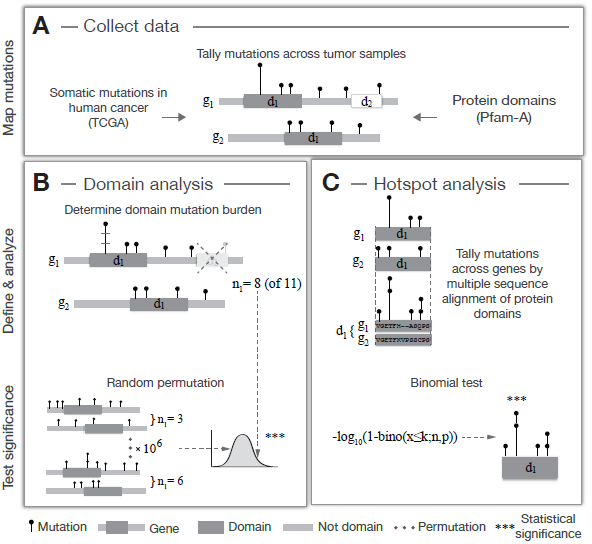
Work flow for analyzing recurrently mutated protein domains in cancer. (**A**) Missense mutation data from recent genomic profiling projects of human cancers (TCGA) are collected and all mutations are tallied across tumor samples and cancer types. Mutations are mapped to protein domains obtained from the Pfam-A database, which contains a manually curated set of highly conserved domain families in the human proteome. Two separate analyses are performed on this data to (**B**) identify domains enriched for missense mutations and (**C**) to detect mutation hotspots in domains through multiple sequence alignment. In the first analysis (**B**), the observed mutation burden (*n_1_*) of a specific domain (*d_1_*) is calculated by counting the total number of mutations in all domain-containing genes (*g_1_ & g_2_*). Mutations in other domains (**e.g.*, d_2_*) are excluded. A permutation test is applied to determine if the observed mutation burden (*n_1_* = 8) is larger than expected by chance. Mutations are randomly shuffled 10^6^ times across each gene separately and the observed mutation count is compared to the distribution of randomly estimated mutation counts. In the second analysis (**C**), domains are aligned across related genes by multiple sequence alignment and mutations are tallied at each residue of the alignment. A binomial test is applied to determine if the number of mutations at a specific residue is significantly different than the number of mutations observed at other residues of the alignment.

We next mapped the mutations to conserved protein domains obtained from the database of protein domain families, Pfam-A version 26.0 (Punta *et al*., 2011) (Fig. 1A). Overall, 4401 of 4758 unique Pfam domains in the human genome were mutated at least once across all samples. The fraction of missense mutations that map to domains (46.7%, 216,676 of 463,842) was consistent across samples and tumor types and was similar to the proportion of the proteome assigned as conserved domains (45.4%, Fig. 2).

**Figure 2:**
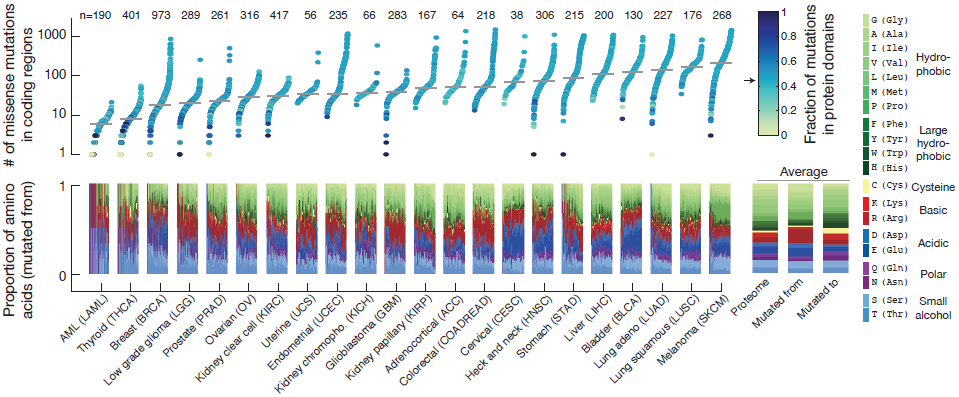
Mutations frequencies across cancer types and the relative proportion of mutated amino acid types. Within each cancer type the individual samples are ordered by the number of missense mutations in the proteome, and the median number of mutations is indicated (grey line). The color code represents the fraction of mutations that map to protein domains, and the arrow indicates the proportion of the proteome assigned as domains (0.454). The lower panel shows the relative proportion of amino acids altered by missense mutations in coding regions in each same (mutated from). The average proportions are displayed on the right where the first bar is the background frequency of amino acids in the proteome, the second bar is the average of all samples (mutated from), and the third bar is the resulting amino acid change (mutated to). The amino acids are color coded by their biochemical properties as indicated. The number of samples in each cancer type is shown at the top. Samples with more than 2000 missense mutations were excluded from the analysis. Note that some amino acid types are disproportionally altered due to mutation biases in specific cancers (Lawrence *et al.,* 2014b; Alexandrov *et al.*, 2013), such as C→G transversions in bladder cancer (BLCA) that disproportionally alter the acidic amino acids aspartic acid (**D**) and glutamic acid (**E**), while C→T transitions in melanoma (SKCM) preferentially affect proline (P).

### Identification of domains with enriched mutation burden

Our first aim was to identify domains that display an increased mutation burden. We defined the domain mutation burden as the total number of missense mutations in a domain, excluding domains only present in only one gene. After tallying mutations across samples, the domain with the highest mutation burden was the protein kinase domain with 7203 mutations in 353 genes (not including genes with tyrosine kinase domains), while the P53 domain present in *TP53, TP63*, and *TP73* had the most mutations when normalizing for the domain length and the size of the domain family (**Supplementary Fig. 3**). To systematically investigate if the mutation burden for a given domain was larger than would be expected by chance, we performed a permutation test that takes into account the number of mutations within and outside of the domain, the domain length, and the length and number of genes in the domain family. To specifically compare domain versus non-domain areas, other domains present in the domain-containing gene family were excluded. Assuming that each mutation is an independent event and that all residues of the protein have an equal chance of being mutated, we randomly reassigned all mutations 10^6^ times across each gene separately and calculated if the observed domain mutation burden was significantly different from the distribution of burdens observed by chance (Fig. 1B).

Using this permutation approach, we identified 14 domains that were significantly enriched for missense mutations within the domain boundaries compared to other areas of the same genes (*p* < 0.05, Bonferroni corrected, Fig. 3 and Table 1). As both the number of gene members per domain (domain family) and the number of mutations per gene varies greatly, we wanted to distinguish between two cases: 1) only a single or a few genes in the domain family contributed to the domain mutation burden, and 2) genes contributed more evenly to the mutations in the domain. We were particularly interested in the latter as mutations in domains contributed by many infrequently mutated genes may represent new functional alterations that would not have been discovered using traditional gene-by-gene approaches. To investigate this, we calculated a entropy score (S̄) that was normalized to the size of the domain family, so a low score indicates that the mutation burden is unevenly distributed between domain-containing genes and a high score indicates that the mutation burden is distributed evenly among the genes in the domainfamily (**see Methods**).

**Figure 3:**
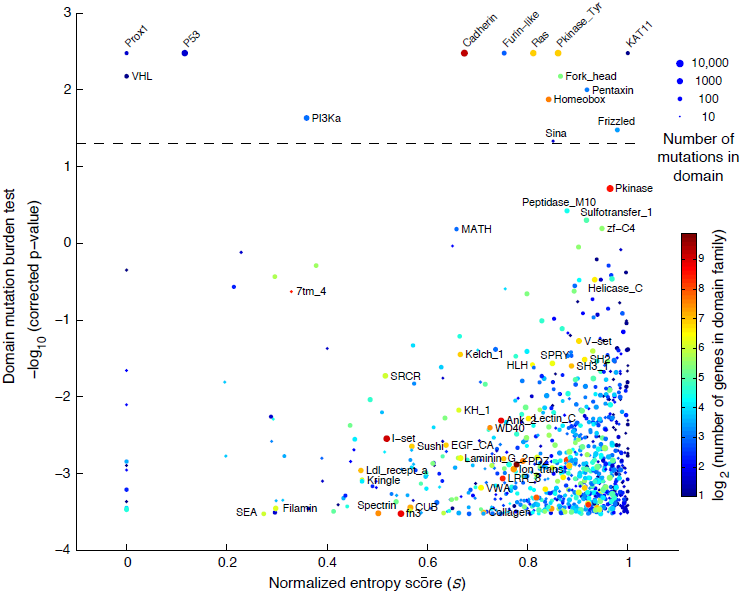
Identification of protein domains enriched for missense mutations. The estimated significance level of the domain mutation burden test is plotted against the domain entropy score (S≃). The domain mutation burden test captures the enrichment of mutations within the domain boundaries compared to non-domain areas of the same genes. The entropy score captures the degree to which individual or multiple genes contribute to the mutations in the domain, where low and high scores indicate that the mutation burden is unevenly or evenly distributed between domain-containing genes, respectively. S̄ is normalized to the highest possible score so that when S̄ = 1 all genes are evenly mutated and when S̄ = 0 then only one gene is mutated. Domains with a significant mutation burden are indicated above the dashed line (*p* < 0.05, Bonferroni corrected). The sizes of the dots reflect the number of mutations in each domain. Domains are color coded by
the number of genes in the domain family.

**Table 1:**
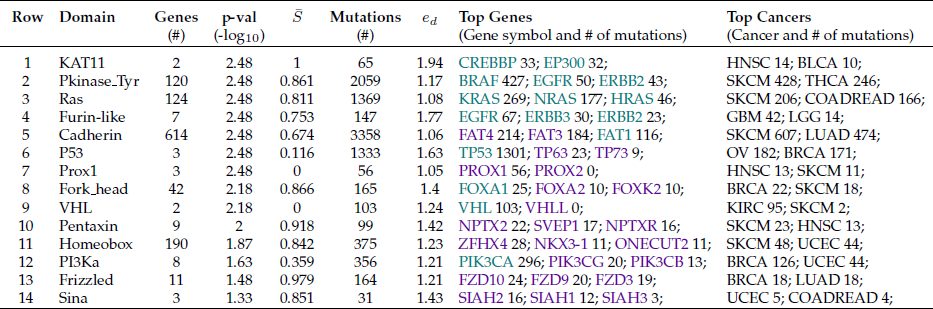
Protein domains significantly enriched for mutations. Domains are listed by their Pfam domain identifiers, the number of genes in the domain family, the Bonferroni-corrected p-value, the entropy score (S̄), the number of mutations in the domain, the mutation enrichment score (*e_d_*) expressed as the ratio of the observed number of domain mutations to the expected number of domain mutations, the genes with the most domain mutations, and the two cancers with most domain mutations. The genes are color coded based on being reported as significantly mutated (green) or not (magenta) in any cancer type in a recent pan-cancer study (Lawrence *et al*., 2014a). The list is sorted by p-value followed by entropy score.

| Row | Domain | Genes (#) | p-val (-log<sub>10</sub>) | $\bar{S}$ | Mutations (#) | $e_d$ | Top Genes (Gene symbol and # of mutations) | Top Cancers (Cancer and # of mutations) |
| --- | --- | --- | --- | --- | --- | --- | --- | --- |
| 1 | KAT11 | 2 | 2.48 | 1 | 65 | 1.94 | CREBBP 33; EP300 32; | HNSC 14; BLCA 10; |
| 2 | Pkinase_Tyr | 120 | 2.48 | 0.861 | 2059 | 1.17 | BRAF 427; EGFR 50; ERBB2 43; | SKCM 428; THCA 246; |
| 3 | Ras | 124 | 2.48 | 0.811 | 1369 | 1.08 | KRAS 269; NRAS 177; HRAS 46; | SKCM 206; COADREAD 166; |
| 4 | Furin-like | 7 | 2.48 | 0.753 | 147 | 1.77 | EGFR 67; ERBB3 30; ERBB2 23; | GBM 42; LGG 14; |
| 5 | Cadherin | 614 | 2.48 | 0.674 | 3358 | 1.06 | FAT4 214; FAT3 184; FAT1 116; | SKCM 607; LUAD 474; |
| 6 | P53 | 3 | 2.48 | 0.116 | 1333 | 1.63 | TP53 1301; TP63 23; TP73 9; | OV 182; BRCA 171; |
| 7 | Prox1 | 3 | 2.48 | 0 | 56 | 1.05 | PROX1 56; PROX2 0; | HNSC 13; SKCM 11; |
| 8 | Fork_head | 42 | 2.18 | 0.866 | 165 | 1.4 | FOXA1 25; FOXA2 10; FOXK2 10; | BRCA 22; SKCM 18; |
| 9 | VHL | 2 | 2.18 | 0 | 103 | 1.24 | VHL 103; VHLL 0; | KIRC 95; SKCM 2; |
| 10 | Pentaxin | 9 | 2 | 0.918 | 99 | 1.42 | NPTX2 22; SVEP1 17; NPTXR 16; | SKCM 23; HNSC 13; |
| 11 | Homeobox | 190 | 1.87 | 0.842 | 375 | 1.23 | ZFHX4 28; NKX3-1 11; ONECUT2 11; | SKCM 48; UCEC 44; |
| 12 | PI3Ka | 8 | 1.63 | 0.359 | 356 | 1.21 | PIK3CA 296; PIK3CG 20; PIK3CB 13; | BRCA 126; UCEC 44; |
| 13 | Frizzled | 11 | 1.48 | 0.979 | 164 | 1.21 | FZD10 24; FZD9 20; FZD3 19; | BRCA 18; LUAD 18; |
| 14 | Sina | 3 | 1.33 | 0.851 | 31 | 1.43 | SIAH2 16; SIAH1 12; SIAH3 3; | UCEC 5; COADREAD 4; |

As expected, we found that the Von Hippel-Lindau (VHL) and the P53 domains were significantly enriched for mutations and had low entropy scores as they were dominated by mutations in the canonical tumor suppressor genes *VHL* and *TP53*, respectively (Fig. 3 and Table 1, row 6 and 9). On the other end of the spectrum, the KAT11 domain encoding the lysine acetyltransferase (KAT) activity of CREBBP and *EP300* was significantly mutated and had a high normalized entropy score with around 30 mutations in each gene (Table 1, row 1). *CREBBP* and *EP300* are transcriptional coactivators that regulate gene expression through acetylation of lysine residues of histones and other transcription factors (Liu *et al*., 2008). In our analysis, head and neck squamous cell carcinoma (HNSC) was the tumor type with most mutations in KAT11 and nearly half fell in the domain (14 of 29) that spans only about 4% of the length of both *CREBBP* and *EP300* (**Supplementary Fig. 4**). Supporting the enrichment of mutations in this domain, inactivating mutations in KAT11 have been associated with oncogenesis in tumor types not part of this analysis, including B-cell lymphoma (Pasqualucci *et al*., 2012; Cerchietti *et al*., 2010; Morin *et al*., 2012) and small-cell lung cancer (Peifer *et al*., 2012). In small-cell lung cancer, *CREBBP* and *EP300* were reported to be deleted in a mutually exclusive fashion (Peifer *et al*., 2012), which often indicates that genes are functionally linked (Ciriello *et al*., 2012). In HNSC, we also found that mutations in *CREBBP* and *EP300* tend to occur in a mutually exclusive pattern (**Supplementary Fig. 4**), indicating a potential function role for both genes in this disease although only one of the two (*EP300*) has previously been linked to HNSC (Lawrence *et al*., 2014a).

Confirming canonical mutation events in cancer, we found mutations clustering in domains of genes involved in receptor tyrosine kinases (RTKs) signaling, including the tyrosine kinase domain itself (Pkinase_Tyr), the furin-like domain involved in RTK aggregation, and downstream signaling through genes with the ras GTPase domain and the phosphatidylinositol 3-kinase (PI3Ka) domain (Table 1, row 2, 3, 4 and 12). These domains have also been reported in other systematic studies of mutations in domains (Yue *et al*., 2010; Nehrt *et al*., 2012), consistent with the fact that the RTK signaling pathways are often high-jacked in cancer (Hanahan & Weinberg, 2011). In a similar manner, we identified multiple domains in genes that have previously been associated with cancer, including the DNA-binding forkhead domain in Fox family transcription factors and the frizzled domain in G protein-coupled receptors of the Wnt signaling pathway. Interestingly, these domains have high entropy scores with a substantial amount of mutations contributed by genes not reported as altered in a recent pan-cancer study (Lawrence *et al*., 2014a) (see color code in Table 1, row 8 and 13). Thus, from the perspective of the structure-function relationship encoded in domains, these are candidate cancer driver genes due to the enrichment of mutations in these functional regions.

We also identified several domain families in which most of the genes had no apparent link to cancer. Such domains include the homeobox domain involved in DNA-binding and the cadherin domain involved in cell adhesion (Table 1, row 5 and 11). As cell-cell adhesion and DNA-binding are critical cellular processes, it is plausible that domain-contained genes involved in these processes are under positive selective pressure in the cancer environment, although it remains to be tested if mutations in these domains are functionally disruptive and may play a critical role in cancer. Several additional domains were found to be enriched for mutations and may potentially be of interest in a cancer context (Table 1).

### Protein domain alignment reveals mutation hotspots across related genes

We next aligned each domain using multiple sequence alignment and tallied mutations across analogous residues of domain-containing genes (Fig. 1C). The goals of this analysis were to identify new domain hotspots with recurrent mutations across functionally related genes and to associate hotspots in well-established cancer genes with rare events in genes not previously linked to cancer. We used a binomial test to determine if a mutation peak at a specific residue was significantly different from other residues in the domain alignment, and we applied the same entropy analysis to investigate the degree to which individual or multiple genes contributed mutations to each hotspot. In total we identified 82 significant hotspots in 42 different domains (**Supplementary Table 1**).

**Figure 4:**
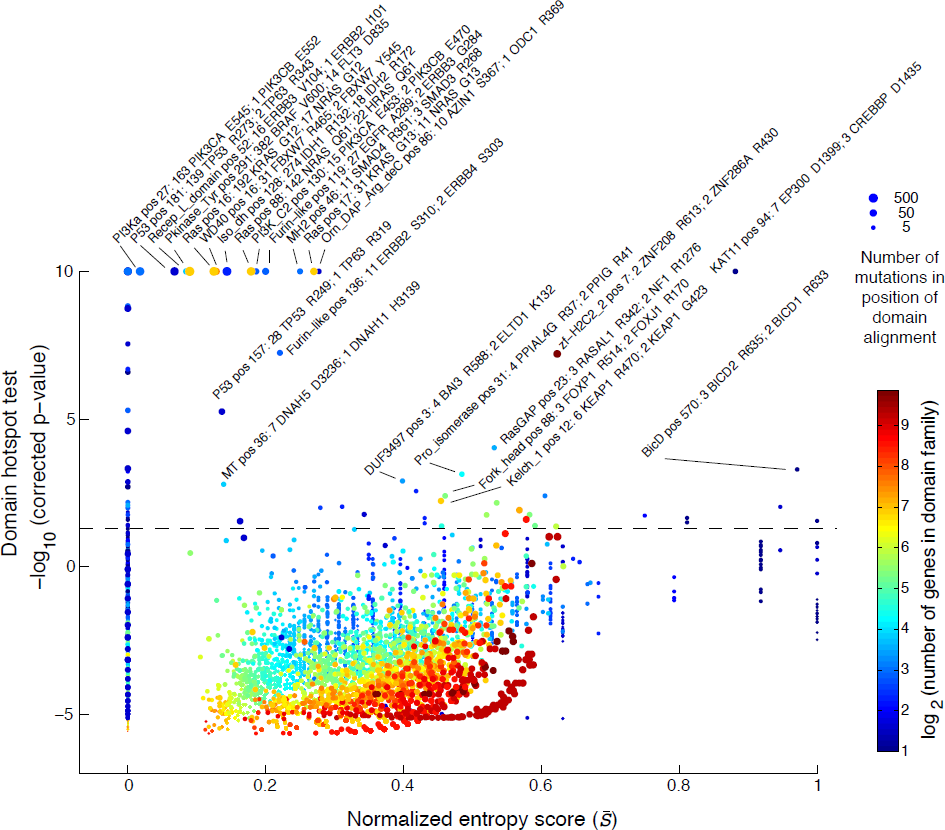
Domain alignment detects mutation hotspots across related genes. The estimated significance level of each mutation hotspot in the domain alignment is plotted against the domain entropy score (S̄), which is described elsewhere. Significant hotspots are indicated above the dashed line (*p* < 0:05, Bonferroni corrected). The maximal significance was set to 10 [-log_10_(p-value)]. Hotspots are named by the Pfam identifiers followed by the position in the domain alignment and the number of mutations in the top two mutated genes. The size of the dots reflects the number of mutations at each residue and the dots are color coded by the number of domaincontaining genes in the genome.

We recapitulated several well-known hotspots in domains where only one gene was mutated such as the P53 and PI3Ka domains with mutations in *TP53* and *PIKC3A*, respectively (entropy⋍0, Fig. 4 and Table 2, row 13, 14, and 17). We also confirmed several known domain-specific hotspots such as the isocitrate/isopropylmalate dehydrogenase domain (Iso_dh) with homologous mutations in *IDH1* (position R132) and *IDH2* (R172) as well as the ras domain with mutations in *KRAS, NRAS*, and *HRAS* at positions G12, G13, and Q61 in the GTP binding region (Table 2, row 3, 7, 8, and 10). Furthermore, we found that well-characterized hotspots in *KIT* D816 in acute myeloid leukemia (AML), *FLT3* D835 in AML, and *BRAF* V600 in thyroid carcinoma and melanoma aligned perfectly in the conserved activation segment of the tyrosine kinase domain (Table 2, row 11). These mutations are known to cause constitutive kinase activity, which promotes cell proliferation independent of normal growth factor control (Hanahan & Weinberg, 2011; Dibb *et al*., 2004). We further superimposed the crystal structures of the three proteins and found that the residues overlap in structure space (**Supplementary Fig. 5**), offering support that the alignment approach captures structurally relevant information. Notably, in the same domain hotspot many singleton mutations in lung adenocarcinoma and lung squamous cell carcinoma mapped to the equivalent position in other RTKs, including *EPHA2* V763 M, *FGFR1 D647 N, PDGFRA* D842H, and three mutations in *EGFR* L861Q. Although these are rare events in lung cancer, this analysis reveals that they likely affect the same activation loop residue and may be therapeutically actionable in a similar manner as the hotspot mutations in *KIT, FLT3*, and *BRAF*. Encouragingly, non-small cell lung cancer patients with *EGFR* L861 mutations have recently shown positive clinical response when treated with *EGFR*-targeted therapy (Wu *et al*., 2011).

**Table 2:**
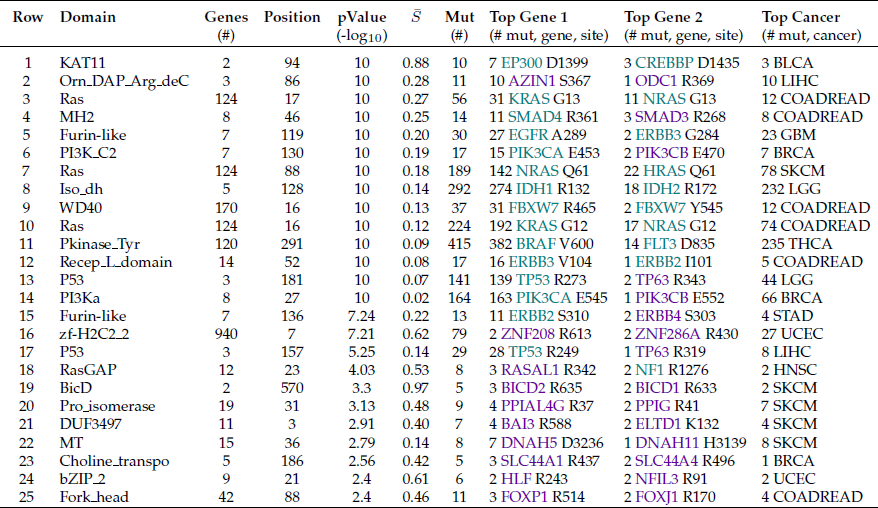
Identified mutation hotspots in protein domains. The detected domain hotspots are listed by their Pfam domain identifiers, the number of genes in the domain family, the position of the hotspot in the domain alignment, the Bonferroni-corrected p-values, the entropy score (S̄), the number of mutations in the hotspot, the two genes with the most mutations in the hotspot, andthe cancer type with most mutations in the hotspot. The genes are color coded based on being reported as significantly mutated (green) or not (magenta) in a recent pan-cancer study (Lawrenceet al., 2014a). The list is sorted by p-value followed by entropy score. Hotspots in domains where only one gene was mutated (*S* = 0) were excluded. All significant domain hotspots (82) areprovided in (**Supplementary Table 1**).

Similar to the previous analysis of entropy in recurrently mutated domains, we were interested in domain hotspots with high entropy scores. Again, the lysine acetylase domain, *KAT11*, was identified with high entropy for a significant hotspot at position 94 of the domain alignment with mutations in *EP300* at D1399 and *CREBBP* at D1435 (Table 2, row 1). These sites are located inthe substrate binding loop of *KAT11* and mutations in these residues affect the structural conformation of the substrate binding loop (Liu *et al*., 2008). Recently, both genes have been implicated in other cancers not analyzed here such as small-cell lung cancer (Peifer *et al*., 2012) and B-cell lymphoma (Pasqualucci *et al*., 2012; Cerchietti *et al*., 2010; Morin *et al*., 2012). Confirming the functional relevance of the identified hotspot, both *EP300* D1399 and *CREBBP* D1435 mutations have been found to reduce lysine acetylase activity in vitro (Peifer *et al*., 2012; Pasqualucci *et al*., 2012; Liu *et al*., 2008). We additionally identified a potential hotspot in *KAT11* at position 105 with mutations in *CREBBP* (R1446) although this hotspot was not significant when correcting for multiple hypothesis testing (*p* = 2.6e^−6^, corrected *p* = 0.59, Fig. 5A). *CREBBP* R1446 is also located within the substrate binding loop (Liu *et al*., 2008) and R1446 mutations have been found in B-cell neoplasms (Pasqualucci *et al*., 2012).

**Figure 5:**
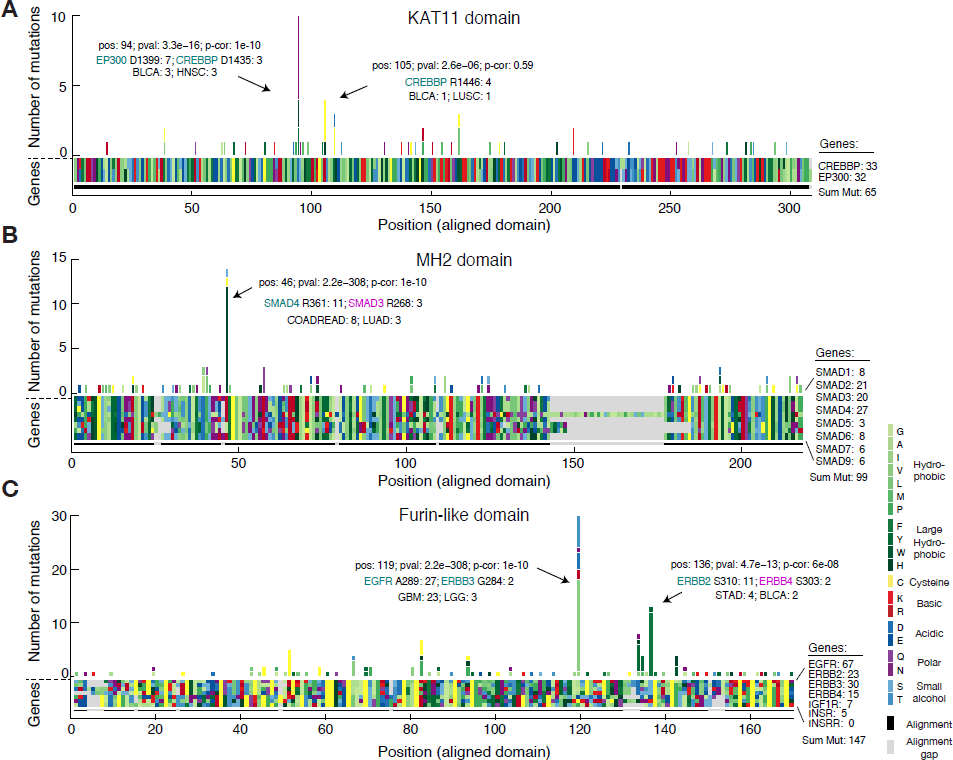
Multiple sequence alignment of domains identifies mutation hotspots and associates rare mutations with known oncogenic hotspots. **(A)** The amino acid sequence alignment of the KAT11 histone acetylate domain in *CREBBP* (position 1342–1648) and *EP300* (position 1306–1612) is represented as a block of two by 308 rectangles. Using the resulting alignment coordinates, missense mutations are tallied across the domains of the two genes. Both amino acids of the alignment (block) and the resulting amino acids due to mutations (histogram) are color coded by their biochemical properties. Alignment gaps are indicated by gray rectangles. Significant hotspots are indicated with position in alignment (pos), p-value (pval), Bonferoni corrected pvalue (p-cor), and number of mutations in top mutated genes and cancer types. Similar plots are shown for the MAD homology 2 (MH2) domain involved in SMAD protein-protein interactions (**B**) and the furin-like domain involved in RTK aggregation and signal activation (**C**).

### Associating rare mutations with known oncogenic hotspots

The MAD homology 2 (MH2) domain is found in *SMAD* genes and mediates interaction between SMAD proteins and their interaction partners through recognition of phosphorylated serine residues (Wu *et al*., 2001). We found the known R361H/C hotspot mutation in *SMAD4* (Shiet al., 1997; Ohtaki *et al*., 2001) aligned with three R268H/C mutations in *SMAD3* (Table 2, row 4). In both proteins these residues are located in the conserved loop/helix region that is directly involved in binding *TGFBR1* (Shi *et al*., 1997). R361C mutations inactivate the tumor suppressor *SMAD4* (Shi *et al*., 1997), and recently, R268C mutations in *SMAD3* were also found to repress *SMAD3*-mediated signaling (Fleming *et al*., 2013), supporting our association of rare arginine mutations in *SMAD3* with known inactivating mutations in *SMAD4*. The majority of the mutations in the hotspot were from colorectal adenocarcinoma samples (COADREAD) and it is known that *SMAD* genes are recurrently mutated in this disease (Fleming *et al*., 2013). Interestingly, we found a tendency towards better survival for patients with hotspot mutations in colorectal cancer although more data is needed to confirm this, to our knowledge, unreported observation (**Supplementary Fig. 6**).

We associated several known hotspots in well-characterized cancer genes with rare but potentially functional mutations in genes not frequently mutated in cancer. For example, we found rare mutations in PIK3_CB at E470 and at E552 in the PI3KC2 domain and PI3Ka domain, respectively, that associated with known recurrent hotspots in PIK3CA (Table 2, row 6 and 14). Furthermore, in the cysteine-rich Furin-like domain, which is involved in receptor aggregation and signaling activation of *ERBB*-family RTKs, we identified several significant hotspots including rare mutations in *ERBB3* (G284 R) and *ERBB2* (A293 V) that aligned with the known activating driver mutations in *EGFR* (A289V/T) in glioblastoma (Lee *et al*., 2006) (Fig. 5C). Recently, one of these mutations, *ERBB3* G284 R, was found to promote tumorigenesis in mice (Jaiswal *et al.*, 2013), suggesting that the singleton *ERBB2 A293 V* mutation found in a melanoma sample could represent an infrequent oncogenic event. We also identified a hotspot at position 137 of the alignment with rare S303 F mutations in *ERBB4* aligning with S310F/Y mutations in *ERBB2*. Interestingly, in a functional analysis of *ERBB2* mutations in lung cancer cell lines, S310F/Y mutations were found to increase *ERBB2* signaling activity, promote tumorigenesis, and enhance sensitivity to *ERBB2* inhibitors in vitro (Greulich *et al*., 2012). Future work will show if analogous mutations in *ERBB4* (S303F) may have similar effects.

### Identification of new hotspots in protein domains

We also identified several additional hotspots in domains with mutations in genes not previously associated with cancer. We detected a hotspot in the prolyl isomerase domain (Pro_fisomerase) with nine mutations distributed between *PPIAL4G* (R37C), *PPIG* (R41C), *PPIA* (R37C), *PPIE* (R173C) and *PPIL2* (I308 F) (Fig. 6A). The Pro_isomerase domain-containing genes catalyze cistrans isomerization of proline imidic peptide bonds and have been implicated in folding, transport, and assembly of proteins (Göthel & Marahiel, 1999). Seven of the nine mutations found in this hotspot were from melanoma samples, and interestingly we found that in melanoma these mutations correlate with significant upregulation of about a dozen genes including the cancer-testis antigens *CTAG2, CTAG1B, CSAG2*, and *CSAG3* (**Supplementary Fig. 7**).

**Figure 6:**
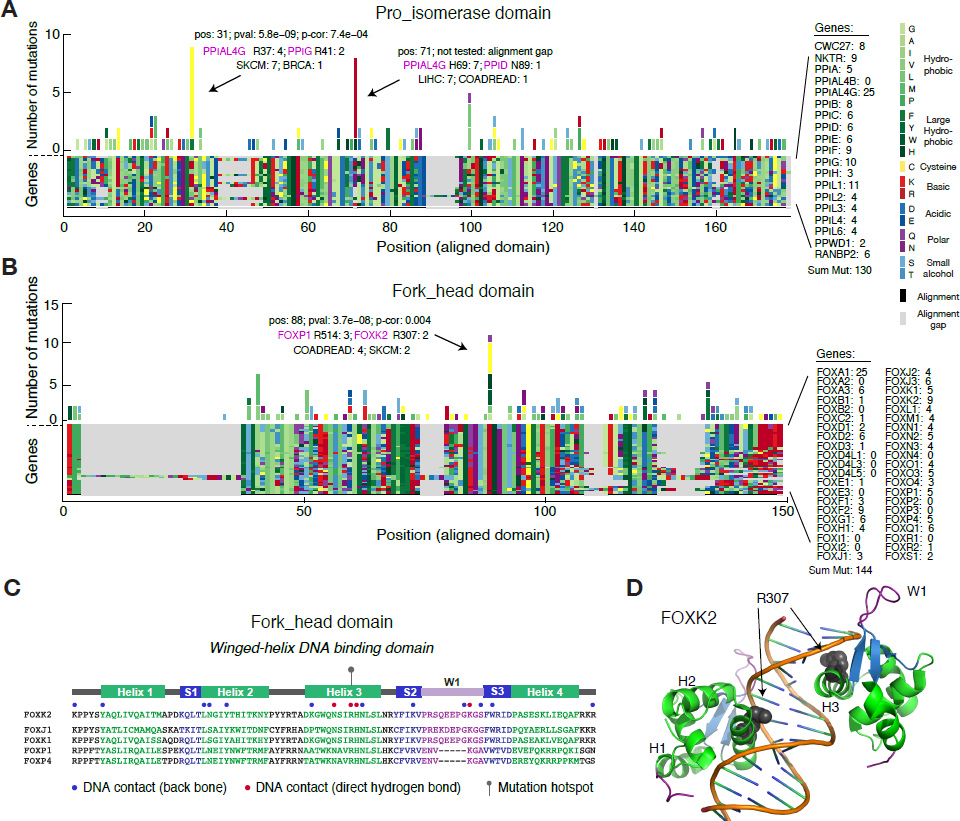
Identification of new hotspots affecting conserved residues of protein domains. Missense mutations are tallied across multiple sequence alignments of genes containing the prolyl isomerase (Pro_isomerase) domain (**A**) and the forkhead domain (**B**). (**C**) Secondary structure of the forkhead domain consisting of four *α*-helices (H1-4), three *β*-sheets (S1-3), and one wing-like loop (W1). Sequences are shown for selected Fox transcription factors that had mutations in the identified hotspot in H3. Of note, the selected genes have a fourth *α*-helix rather than the canonical second wing-like loop found in other Fox genes. (**D**) Ribbon drawing of the crystal structure of two *FOXK2* forkhead domains binding to a 16-bp DNA duplex containing a promoter sequence (pdb ID: 2C6Y) (Tsai *et al.*, 2006). The R307 residue that we identified as mutated in the hotspot is shown with spheres.

The forkhead domain mediates DNA binding of forkhead box (Fox) transcription factors and encodes a conserved “winged helix” structure comprising three α-helices and three β-sheets flanked by one or two “wing”-like loops (Carlsson & Mahlapuu, 2002). In the forkhead domain, we identified a hotspot with 11 mutations distributed between *FOXP1* (R514C/H), *FOXK2* (R307C/H), *FOXK1* (R354 W), *FOXJ1* (R170G/L), and *FOXP4* (R516C) in several different cancer types (Fig. 6B). The identified hotspot was located in the third α-helix (H3), which exhibits a high degree of sequence homology across Fox proteins and binds to the major groove of DNA targets. Specifically, the arginine residue that we found mutated forms direct hydrogen bonding with DNA in both in *FOXP* and *FOXK* family transcription factors (Wu *et al*., 2006; Stroud *et al*., 2006; Chu *et al*., 2011) (Tsai *et al*., 2006) (Fig. 6C, D). Furthermore, experimental R307A substitution in *FOXK2* abolishes DNA binding (Tsai *et al*., 2006), suggesting that the identified arginine mutations may play an important role in cancer by inhibiting DNA-binding of *FOXP, FOXK*, and related Fox transcription factors.

We identified several other domain hotspots of potential interest such as a hotspot in the ras-GAP GTPase activating domain with mutations in the tumor suppressors *NF1, RASA1*, and *RASAL1* (**Supplementary Fig. 8**) and a hotspot in the kelch motif (Kelch_1 domain) with mutations in KEAP1 and KLHL4 (**Supplementary Fig. 9**). The potential biological consequence of these mutations remains to be elucidated. Many additional domain hotspots were identified and we make all analysis of hotspots in protein domains available via an interactive web-service at http://www.mutationaligner.org.

## DISCUSSION

In this work we used protein domains, rather than individual genes, as the basis for the discovery of cancer-relevant alterations. By coupling the observation of mutations across common members of a domain family together, we identified domains enriched for mutations as well as mutation hotspots within these domains. Many of these domain mutations were contributed by groups of relatively rare mutations which that otherwise have been considered as spurious passengers. We further associated putative function with infrequent mutations as we identified hotspots where rare and well-characterized mutations affected analogous residues in the domain alignments. Based on the structure-function relationship encoded in domains, these rare mutation events may potentially be therapeutically actionable in cases where drugs have been developed to target related genes sharing the mutated domain. Finally, we identified entirely new hotspots in domains with mutations in genes not previously associated with cancer, hereby nominating new potential cancer-related genes for further investigation.

The fundamental assumptions underlying this work are that mutations at analogous sites of a domain family have a common effect and that recurrent mutations in domains are likely associated with cancer. As many protein domains have been functionally characterized, one of the strengths of our approach is that such knowledge can provide mechanistic insight into the potential effect of alterations. For example, we confirmed that canonical signaling domains present in large gene families are enriched for mutations (*e.g.*, tyrosine kinase and ras domains), reflecting the fact that a range of genes involved in mitogenic signaling are often highjacked in cancer (Hanahan & Weinberg, 2011). We also identified less characterized domains that were significantly altered, such as the homeobox domain of DNA-binding genes and the cadherin domain of cell adhesion genes. While the individual domain-containing genes are not recurrently altered, these new findings suggest that mutations in the DNA-binding and cell adhesion machinery as general phenomena may confer a selective advantage in cancer. Similarly, in the prolyl isomerase domain, we identified a new hotspot in melanoma with arginine to cysteine mutations in *PPIA, PPIG*, and *PPIAL4 G*. We also identified a new hotspot in the forkhead domain, where the crucial DNA-contacting arginine residue in the third helix of the forkhead-encoded winged-helix structure was mutated in several *FOXP* and *FOXK* family transcription factors. We speculate that this is a novel inactivating oncogenic event, although more research is needed to elucidate this.

Relatively few genes are recurrently mutated in cancer and the majority of somatic mutations are observed in infrequently mutated genes (Stephens *et al*., 2013; Garraway & Lander, 2013). As above, we can use the biological knowledge associated with protein domains to help interpret the consequences of rare mutations in well-characterized domain families. For example, in the furin-like domain, we associated known oncogenic *ERRB2 S310F/Y* mutations with uncharacterized *ERBB4 S303 F* mutations. In both genes, a small amino acid with an alcohol side chain (S) is mutated to a large hydrophobic amino acid (F or Y) in a conserved region involved in receptor interaction and signal activation. S310F/Y mutations in *ERBB2* are tumorigenic in vitro and it has been speculated that S310F/Y mutations promote hydrophobic interactions and receptor dimerization resulting in receptor activation (Greulich *et al*., 2012). The same report also found that S310 F mutations sensitize cell lines to the RTK inhibitors neratinib, afatinib, and lapatinib. We identified *ERBB4 S303 F* mutations in breast and endometrial cancers and predict that *ERBB4* S303 F mutations are gain-of-function mutations that increases sensitivity to small-molecule inhibition and therefore represents a rare but druggable oncogenic event. Additionally, in the MH2 domain we associated known loss-of-function mutations *SMAD4* (R361H/C) in colorectal and lung cancers with rare but potentially functional mutations in *SMAD3* (R268H/C) in the same cancers. These examples illustrate how the association of mutations in infrequently altered genes with known mutations can provide high-confidence predictions and hypotheses for further experimental testing.

By analyzing mutations in a set of functionally related genes, we introduce the risk of detecting false positive hotspots where passenger mutations are aggregated across domain-containing genes. The detection of spurious domain hotspots could be exacerbated by mutation biases that alter amino acids disproportionally (Fig. 2). For example, arginine is the most frequently altered amino acid as it has four codons containing CG dinucleotides, which are frequently subject to C→T transitions due to deamination of methylated cytosine to thymine. In several domains, such as the tetramerisation (K_tetra) domain of potassium channel proteins (**Supplementary Table 1**) and the zf-H2C2_2 domain of zinc finger proteins (Table 2, row 16), we identify significant hotspots where arginine mutations align across a large set of genes in the domain family. Although the frequency of such hotspots may be driven by the arginine mutation bias, the mutations themselves may nevertheless be functional. Thus, we do not penalize for amino acid mutation biases in the detection of hotspots. However, we do provide relevant information about the potential biases by calculating a normalized hotspot z-score that takes into account the relative amino acid mutation frequency in each cancer type (**Supplementary Table 1** and Methods).

Future work will aim to refine the analysis of mutations in domains and expanding the scope of our analysis to other functional elements in genes. Our focus here was on somatic missense mutations, but this requirement may be relaxed to include germline mutations or other somatic alterations (*e.g.*, truncating mutations and small in-frame insertions and deletions). Importantly, truncating mutations may not necessarily be localized to the domain regions as they affect the entire gene, which is why we excluded them from our current work. An additional extension of our work would be to implement a sliding window for peak detection of clusters of mutations in domain alignments. However, our own observations suggest that mutation hotspots are largely localized to single residues. Structural information can also be used to analyze the proximity of mutated residues in 3D space. Other types of regulatory protein motifs can be analyzed, including short linear motifs that guide protein phosphorylation by kinases, which has previously been shown to be enriched for mutations in cancer genes (Reimand & Bader, 2013; Reimand *et al*., 2013). Finally, assessment of the functional impact of mutations using structural information and evolutionary sequence conservation, for example as applied in our mutation assessor method (Reva *et al*., 2011), can be incorporated to provide additional insight into the potential role of mutations in cancer.

As more data become available, integrative approaches combining mutation evidence across multiple scales such as genes, domains, and signaling pathways will be needed to improve the computational pipelines for variant function prediction. To make the results of our analysis useful and relevant to the community at large, we have made all findings available through an interactive web service (http://www.mutationaligner.org) that will be continuously updated as new tumor samples are genomically profiled.

## EXPERIMENTAL PROCEDURES

### Mutation data and data preprocessing

All TCGA mutation data were obtained in MAF file format from the cBioPortal for cancer genomics data (Cerami *et al*., 2012; Gao *et al*., 2013). To filter out mutations in low expressed genes, which has been shown to be associated with mutation biases (Lawrence *et al*., 2014b), mRNA sequencing data in the form of normalized RSEM values were obtained from the same data portal. Within each tumor type, we determined the mean RSEM value for each gene and mutations in genes with a mean RSEM value of less than 10 were excluded from the analysis. To filter out ultra-mutated cancers, samples with more than 2,000 non-silent mutations were disregarded. The TCGA tumor types analyzed were: Acute myeloid leukemia (LAML), Adrenocortical carcinoma (ACC), Bladder urothelial carcinoma (BLCA), Brain lower grade glioma (LGG), Breast invasive carcinoma (BRCA), Cervical squamous cell carcinoma and endocervical adenocarcinoma (CESC), Colorectal adenocarcinoma (COADREAD), Glioblastoma multiforme (GBM), Head and neck squamous cell carcinoma (HNSC), Kidney chromophobe (KICH), Kidney renal clear cell carcinoma (KIRC), Kidney renal papillary cell carcinoma (KIRP), Liver hepatocellular carcinoma (LIHC), Lung adenocarcinoma (LUAD), Lung squamous cell carcinoma (LUSC), Ovarian serous cystadenocarcinoma (OV), Prostate adenocarcinoma (PRAD), Skin cutaneous melanoma (SKCM), Stomach adenocarcinoma (STAD), Thyroid carcinoma (THCA), Uterine carcinosarcoma (UCS), Uterine corpus endometrial carcinoma (UCEC).

### Pfam domains and mapping mutations to protein domains

The Pfam-A data base of domains in the human proteome (version 26) as well as all human protein sequences were downloaded from the Pfam ftp server (pfam26.9606.tsv, http://ftp://ftp.ebi.ac.uk/pub/databases/Pfam). To include only high confidence domain calls, domains with an expectancy value (e-value) larger than 1e^−5^ were excluded. Mapping entries between MAF files and Pfam domains was performed using Uniprot accession numbers using the MAF ONCOTATOR_UNIPROT_ACCESSION_BEST_EFFECT field. In cases where the MAF entries did not have Uniprot accession numbers, the biomart webservice (http://www.ensembl.org/biomart/) was used to map between HGNC gene symbols and Uniprot accession numbers. The protein domain coordinates from the Pfam-A database were then matched to the MAF entries to determine if the mutations fell within or outside the boundaries of the protein domains using the MAF ONCOTATOR_PROTEIN_CHANGE_BEST_EFFECT field. MAF entries for which the mutated protein position and amino acid identity did not match with the corresponding amino acid identity in the protein sequences were excluded from the analysis. Furthermore, we excluded MAF entries where the mutated protein position was larger than length of the protein sequence.

### Identification of domains with enriched mutation burden

For each domain we tallied the number of missense mutations falling (1) within the domain boundary, and compared it to (2) outside of the boundaries of all other domains in the gene, effectively excluding other domains than the domain in question. To assess if the mutation burden of the domain was larger than would be expected by chance, we implemented a permutation test. The permutation test compared the observed mutation burden of the domain to the distribution of burdens generated by randomly distributing mutations across genes containing the domain. To generate this distribution, we repeated the following process for each permutation *i*:

1. For each gene *g* in the domain family, count the total number of observed mutations in the gene (both within and outside of the domain). Define this quantity to be *n_g_*.
2. For each gene *g*, randomly redistribute ng mutations across the gene, allowing for multiple mutations to fall at the same amino acid residue.
3. Count the total number of mutations which fall within the domain boundaries across all genes. Define this quantity to be *m_i_*, the mutation burden of the domain in permutation *i*.

To calculate a p-value for the observed mutation burden of the domain, we compared the true mutation burden *m_d_* derived from the data to the distribution of *m_i_*. The p-value was defined to be the proportion of permutations with mutation burden greater than or equal to the observed mutation burden.

Note that by treating each gene separately and summing over the outcome of randomly distributed mutations in each gene, we are able to account for gene-to-gene variation in mutation rate (*e.g.*, variation associated with replication timing (Lawrence *et al*., 2014b) as well as differences in gene length and the proportion of each gene occupied by domains).

Domains with less than 25 mutations across all cancer types were excluded in the permutation analysis to avoid spurious results due to low mutation counts. Furthermore, to ensure proper random redistribution of mutations across genes and their domains, we omitted domains where the fraction of amino acids assigned as domains was larger than 75% of the all amino acids in the domain-containing proteins.

### Domain mutation enrichment score

To calculate an enrichment score of mutations in the domain (*e_d_*), we compared the observed domain mutation burden (*m_d_*) to the expected domain mutation burden (*m_e_*). We calculated *m_e_* based on the total number of mutations observed (*n_g_*) and the fraction of amino acids assigned as domains compared to total length of all genes in the domain family (*f_d_*):

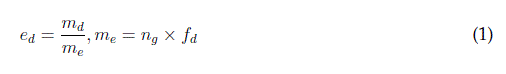

### Multiple sequence alignment of protein domains

The domain amino acid sequences were obtained as substrings from the protein sequences and aligned across domain-containing genes using the MathWorks multialign package with BLOSUM80 as scoring matrix and default parameters. For aligning domains present in only two genes, the Needleman-Wunsch algorithm was by applied using the MathWorks nwalign package with default parameters. After alignment of domains, missense mutations were tallied across analogous residues of domaincontaining genes using the coordinates of the multiple sequence alignment. Residues with alignment gaps in more than 75% of the sequences were excluded from the domain hotspot analysis.

### Identification of mutation hotspots within domain alignments

To identify putative hotspots for mutations within domains, we used as a null model the case of mutations falling with equal likelihood at all sites within a domain. Following multiple sequence alignment of all genes within a domain family, we tallied the number of observed mutations within the domain. We assumed that, for a particular residue to be called a putative hotspot, more mutations must fall on that residue than would be expected by chance if mutations were randomly distributed throughout the body of the domain. Assuming that each mutation falls at a random site along the domain body, the frequency of mutations at any particular residue follows a binomial distribution:

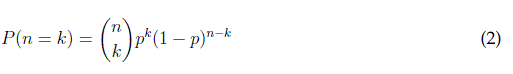

where *n* is the total number sof mutations in the domain, *k* is the number of mutations falling at a particular residue, and *p* is the probability of any individual mutation falling at a particular residue, and *P*(*n* = *k*) is precisely the probability of observing *k* mutations at a single residue, assuming that *n* mutations were observed across the entire domain. Because our null model assumes an equal likelihood of mutations at any residue, 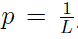, where *L* is the length of the domain.

Thus, to assign a probability to the observation of k mutations falling at a particular site by change (*i.e.*, a p-value), we calculate the probability of at least *k* mutations falling at a particular site from our null model

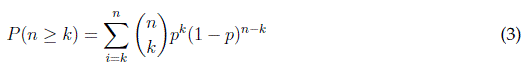

To correct for multiple hypothesis testing, p-values for all considered hotspots (aligned domain residues with more than two mutations) were adjusted using the Bonferroni correction method.

### Calculating z-scores for mutation counts

To provide an estimate whether the number of mutations at a given mutation hotspot (mutation count) is different from the mutation counts at other residues of the domain alignment, we calculated a z-score for each position of the alignment. Assuming a normal distribution, the z-score expresses the number of standard deviations a given hotspot is above the mean of the distribution of mutations counts observed in the alignment. Additionally, we calculated a normalized z-score that takes into account the biases in amino acid mutation frequencies observed in different cancers by calculating the z-score based on a normalized mutation count (*ĉ*):

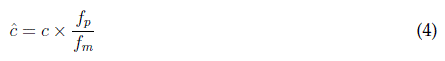

where *c* is the observed mutation count, *f_p_* is the background frequency of a given amino acid in the proteome, and *f_m_* is the observed mutation frequency of a given amino acid in a specific cancer type.

### Entropy calculations

To assess how uniformly the mutations in a specific domain are spread across the genes containing such domain, we rely on the notion of Shannons information entropy. The information entropy *S* of a discrete probability distribution *P(x)* is defined to be

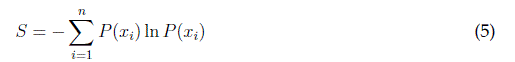

where *P*(*x_i_*) is the probability of the *i^th^* value of *x*. The entropy is maximal when *P*(*x*) is uniform, *i.e.*, each value of *x* is equally probable (*S_max_* = ln *n*), and minimal when *P*(*x*) is equal to 1 for a single value of *x* (*S_min_* = 0). In order to facilitate the comparison of entropy values for vectors of different dimension (*e.g.*, domain families with different numbers of constituent genes), we use a normalized entropy measure *S̄* defined as

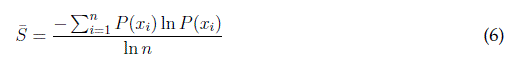

where *n* is the dimension of the vector *x*.

## Author contributions

M.L.M. and E.R. designed analysis. N.P.G. designed and developed the web-site. M.L.M., E.R., N.P.G, B.A.A., A.K., J.G., and N.S. analyzed data. M.L.M. conceived and developed the concept. M.L.M. and C.S. managed the project. All authors contributed to discussions and editing of the manuscript.

## Acknowledgments

We acknowledge C. Kandoth for technical assistance. C. Carmona-Fontaine and A. Hanrahan for helpful discussions. V.A. Pedicord for helpful comments on the manuscript. This work was funded in part by the National Cancer Institute Cancer Genome Atlas grant (U24 CA143840).

